# Purification of mitochondria from skeletal muscle tissue for transcriptomic analyses reveals localisation of nuclear-encoded non-coding RNAs

**DOI:** 10.1101/2022.04.27.489477

**Authors:** Jessica Silver, Adam J. Trewin, Stella Loke, Larry Croft, Mark Ziemann, Megan Soria, Hayley Dillon, Søren Nielsen, Séverine Lamon, Glenn. D. Wadley

**Author notes:** **Corresponding author:** Prof. Glenn Wadley Deakin University Institute for Physical Activity and Nutrition, School of Exercise and Nutrition Sciences, Australia 3125. (co-first authors).

## Abstract

Mitochondria are central to cellular function, particularly in metabolically active tissues such as skeletal muscle. Nuclear-encoded RNAs typically localise within the nucleus and cytosol but a small population may also translocate to subcellular compartments such as mitochondria. We aimed to investigate the nuclear-encoded RNAs that localise within the mitochondria of skeletal muscle cells and tissue. Intact mitochondria were isolated via immunoprecipitation (IP) followed by enzymatic treatments (RNase-A and proteinase-K) optimised to remove transcripts located exterior to mitochondria, making it amenable for high-throughput transcriptomic sequencing. Small-RNA sequencing libraries were successfully constructed from as little as 1.8 ng mitochondrial RNA input. Small-RNA sequencing of mitochondria from rat myoblasts revealed the enrichment of over 200 miRNAs. Whole-transcriptome RNA sequencing of enzymatically-purified mitochondria isolated by IP from skeletal muscle tissue showed a striking similarity in the degree of purity compared to mitoplast preparations which lack an outer mitochondrial membrane. In summary, we describe a novel, powerful sequencing approach applicable to animal and human tissues and cells that can facilitate the discovery of nuclear-encoded RNA transcripts localised within skeletal muscle mitochondria.

## INTRODUCTION

Mitochondria are subcellular organelles essential for numerous cellular processes including energy metabolism, signalling, ion and redox homeostasis as well as regulation of apoptosis (Spinelli and Haigis, 2018). Mitochondria are comprised of proteins encoded by both the nuclear and mitochondrial genomes, but nuclear-encoded RNAs are not usually localised within mitochondria. A small number of specific nuclear-encoded non-coding RNAs (ncRNAs), such as microRNAs (miRNAs) (Barrey et al., 2011, Sripada et al., 2012, Das et al., 2012) and possibly long non-coding RNAs (lncRNAs) (Noh et al., 2016) can however be imported into mitochondria (Silver et al., 2018). This may occur via the polynucleotide phosphorylase (Wang et al., 2010) and other putative transport mechanisms (reviewed in Silver et al., 2018). Because ncRNAs have the ability to regulate various transcriptional and translational processes relating to mitochondrial function, it is of interest to determine which of these may localise within mitochondria in order to understand the nature of their potential regulatory actions.

Investigations into mitochondria-localised nuclear-encoded RNAs have been constrained, first, by the technical challenges of obtaining pure mitochondrial preparations free of confounding RNAs peripheral to isolated mitochondria and, second, by the significantly lower RNA yields from subcellular fractions when compared to whole tissue homogenates. Early studies providing evidence for the localisation of ncRNAs within mitochondria stem from micro-array analysis and were routinely validated by single qPCR assays or *in situ* hybridisation (Barrey et al., 2011, Sripada et al., 2012, Das et al., 2012). However, the overall population of ncRNAs that localise within mitochondria and their regulatory actions, particularly within metabolically active tissues remains largely unknown.

Skeletal muscle constitutes ∼40% of human body mass and plays a central role in whole body metabolism (Zurlo et al., 1990). To meet these dynamic bioenergetic demands, skeletal muscle is densely populated with mitochondria (Russell et al., 2014). Thus, gaining a greater understanding of the RNA profile within muscle mitochondria may yield important insights into the cellular biology and aetiology of diverse metabolic diseases characterised by mitochondrial dysfunction, such as diabetes mellitus and chronic myopathies (Silver et al., 2018). Here, we report a method for isolating and optimising the enzymatic purification of mitochondria from rodent muscle cells or tissues. We provide direct evidence for the purity of this mitochondrial fraction, making it amenable for downstream high-throughput analyses. We also describe methods for small-RNA and whole transcriptome sequencing of mitochondria isolated from cultured myoblasts as well as skeletal muscle tissue samples from animals and humans.

## RESULTS

### Protease purification of isolated mitochondria from myoblasts

We first isolated mitochondria from cultured L6 rat myoblasts using an antibody against an outer mitochondrial membrane protein (TOMM22) conjugated to magnetic beads allowing for precipitation of intact isolated mitochondria (Figure 1). Immunoblot of the post-precipitation mitochondrial fraction revealed that a small amount of α-tubulin and Lamin B1 (used as protein markers of the cytosol and nucleus, respectively) were present in the isolated mitochondria fraction (Figure 2, lane 1). A portion of the isolated mitochondria was incubated with the serine protease, proteinase-K, to digest proteins not located within the mitochondrial intermembrane space (IMS) or matrix compartments. This led to depletion of α-tubulin and Lamin B1, in addition to a marker of the outer mitochondrial membrane (TOMM20), while intermembrane space (cytochrome-*c*) and matrix protein markers (citrate synthase) were protected from digestion (Figure 2, lane 2). All proteins probed were susceptible to proteinase-K when mitochondrial membranes were dissolved in the presence of the detergent triton-X (Figure 2, lane 3). Together, this demonstrates that mitochondrial isolation by TOMM22-immunoprecipitation followed by enzymatic treatment yields a purified, intact mitochondrial fraction suitable for downstream protein analyses. Based on this, we hypothesised that an analogous approach utilising a ribonuclease could be used to remove RNA located outside of isolated mitochondria.

**Figure 1.**
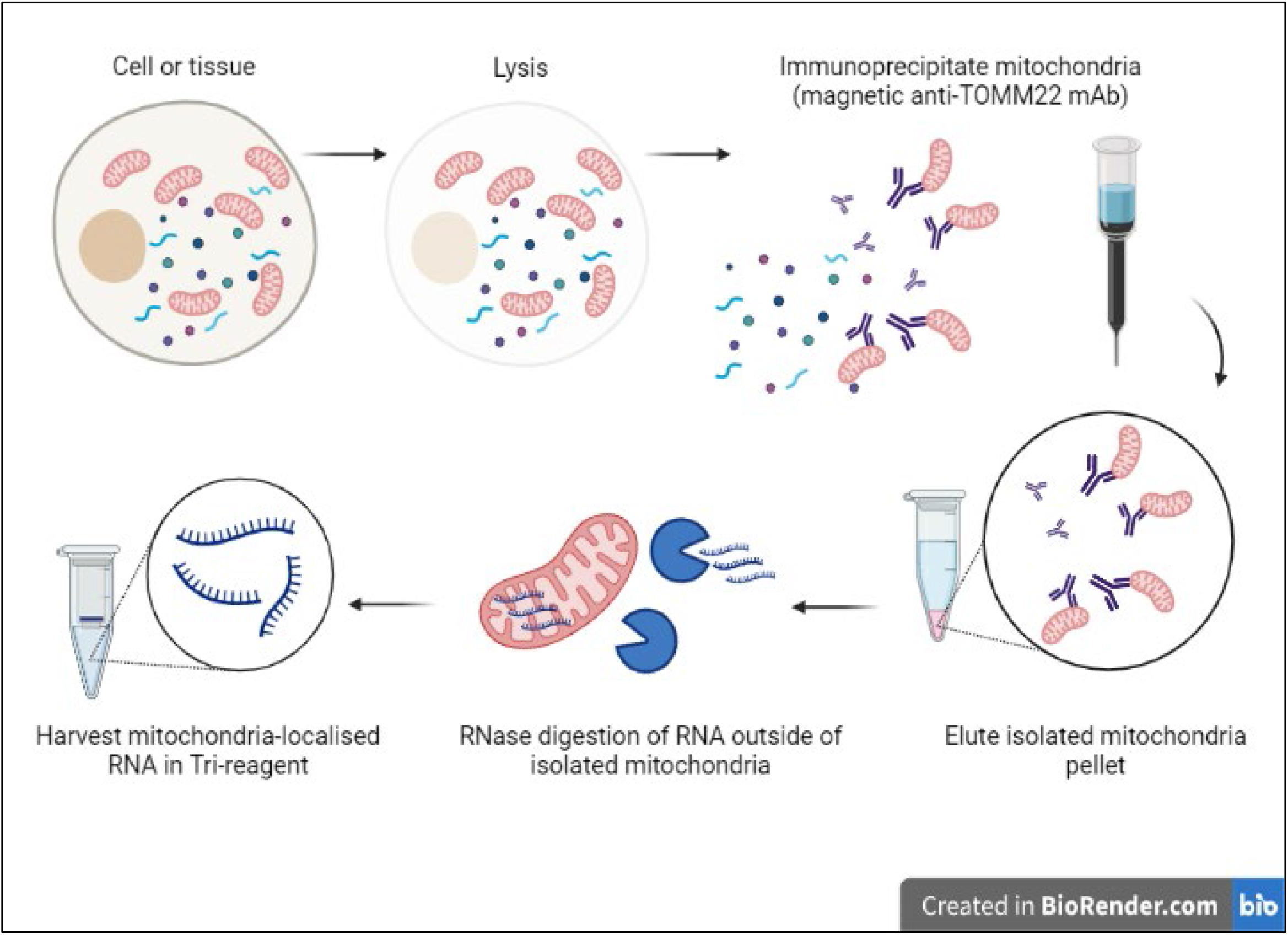
Overview of TOMM22 immunoprecipitation based method for mitochondrial isolation.

**Figure 2.**
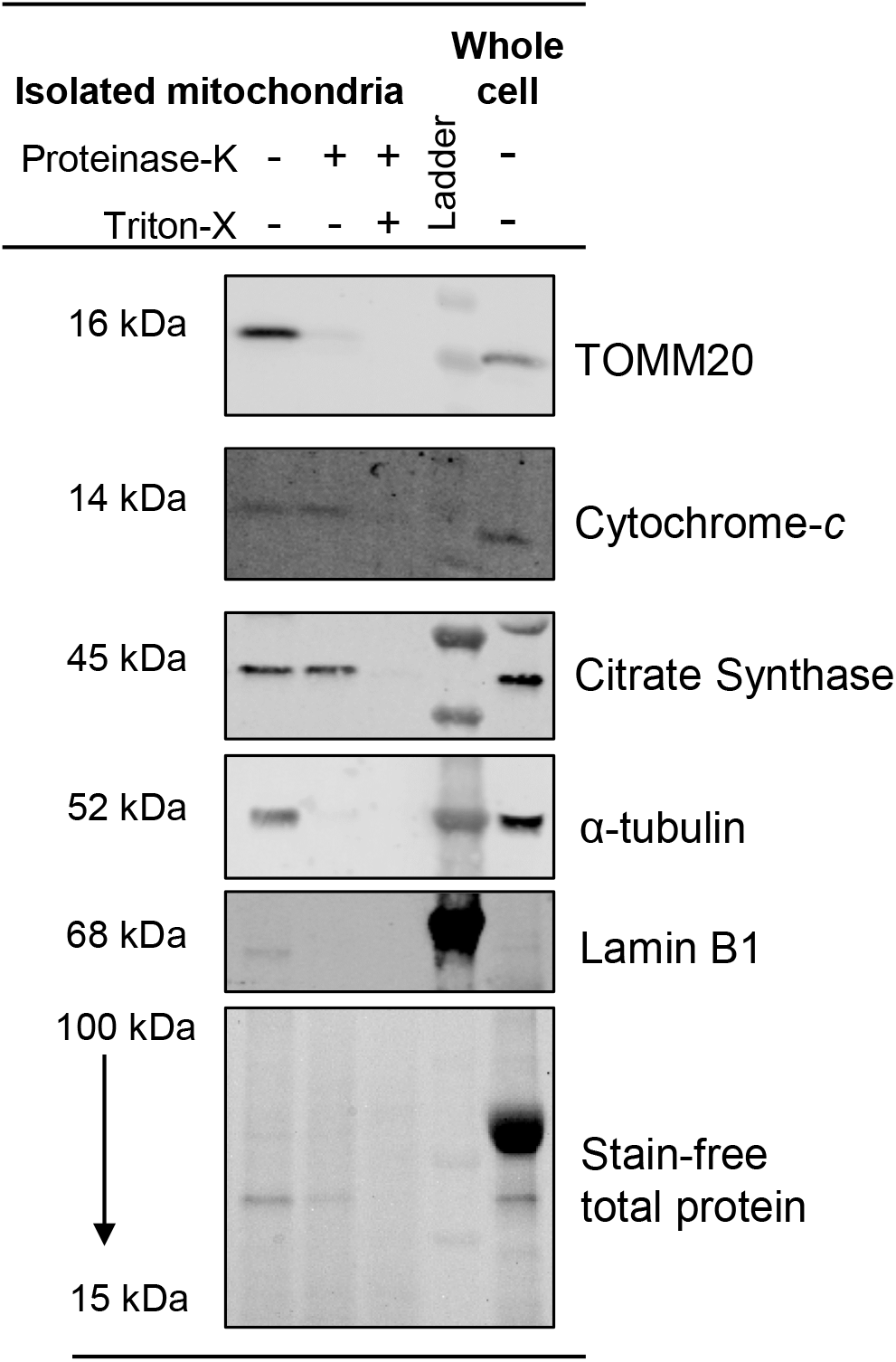
Protease treatment of mitochondria isolated from L6 myocytes. Isolated mitochondria were incubated with or without proteinase-K (20 µg/mL) and detergent (1% triton-X). Mitochondria (0.2 µg total protein) and whole cell lysate (12 µg total protein) were resolved via SDS-PAGE and probed for marker proteins of subcellular compartments: TOMM20 (OMM), Cytochrome-c (IMS), citrate synthase (matrix), α-tubulin (cytosol), lamin B1 (nucleus). Blots are representative of *n*=2 technical replicates from a single mitochondrial isolation preparation.

### Ribonuclease purification of isolated mitochondria from myoblasts

Next, mitochondria isolated from L6 rat myoblasts were exposed to a range of concentrations of RNase-A, an endonuclease that cleaves phosphodiester bonds after pyrimidine residues in the RNA molecule (Witzel, 1963). RNase-A-treated isolated mitochondria were assessed by RT-qPCR for abundance of mtDNA-encoded mRNA (*Mt-co3*) localised exclusively within the mitochondrial matrix compartment, and a nuclear-encoded mRNA (*Cox4i1*) that localises outside mitochondria (Suissa and Schatz, 1982). We found that *Mt-co3* was protected from degradation at RNase concentrations up to at least 100 µg/mL, whereas partial *Cox4i1* degradation occurred at 10 µg/mL and was mostly degraded at 100 µg/mL (Figure 3). In the presence of detergent to disrupt mitochondrial membranes, 1 µg/mL RNase-A was sufficient to digest all *Cox4i1* and *Mt-co3*, as well as an exogenous mRNA spike-in control (Figure 3). This suggests that a certain proportion of cytosolic *Cox4i1* is protected from digestion at lower RNase-A concentrations, potentially as a result of being bound to ribosomes at the outer mitochondrial membrane (Gold et al., 2017). Taken together, these data suggest that TOMM22-IP mitochondrial isolation followed by RNase-A treatment is suitable for downstream mitochondrial RNA analyses provided the RNase-A concentration is sufficient to remove transcripts not located within the mitochondrial membranes.

**Figure 3.**
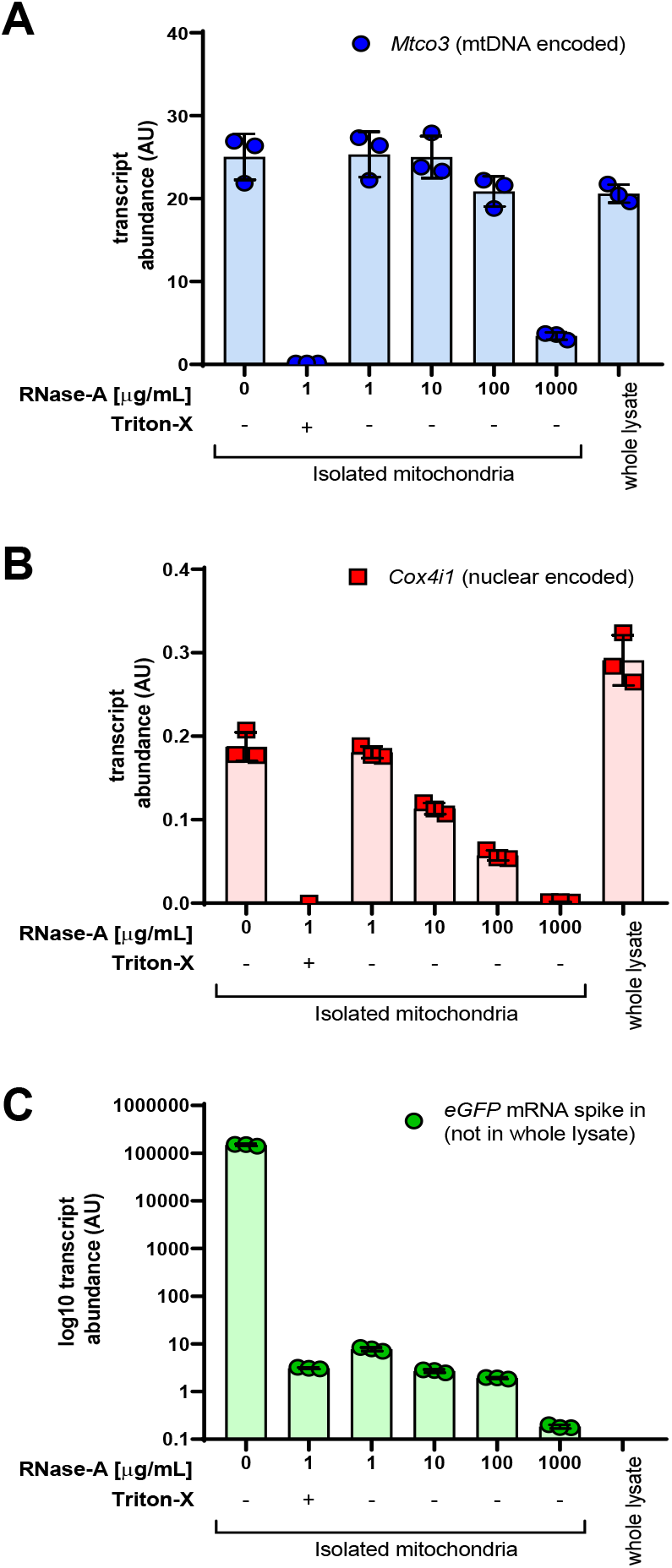
RNase-A treatment of mitochondria isolated from L6 myocytes. Isolated mitochondria were incubated with or without RNase-A (1-1000 µg/mL) in the presence or absence of detergent (1% triton-X). The levels of representative mtDNA (*Mtco3*) and nuclear (*Cox4i1*) encoded genes were then assessed by RT-qPCR in addition to an exogenous mRNA spike-in control (*eGFP*) added to isolated mitochondria (but not to the whole cell lysate). Data are mean(SD) for *N*=3 technical replicates from a single mitochondrial isolation preparation.

### Small RNA sequencing on purified isolated mitochondria from myoblasts reveals mitochondria localised miRNAs

We next aimed to optimize a protocol to perform transcriptomic analysis of the small RNAs (sRNA) contained within isolated mitochondria from L6 rat myoblasts. We first prepared sRNA libraries from 120 ng and 60 ng mitochondrial RNA with 0.5X adapter dilutions (NEBNext 3’ and 5’ SR Adaptors, and RT primer for Ilumina), in line with the standard manufacturer protocol (NEBNext Multiplex Small RNA Library Prep Set (New England Biolabs). The standard protocol produced large amounts of adapter-dimer and was not sufficient to ligate adapters to the target miRNAs within the sample (Supplementary Figure 1 A, B). We next decreased the amount of the 3’ and 5’ adapters to investigate if a lower molar ratio improved miRNA ligation efficiency. Both the modified-0.3X (Supplementary Figure 1C) and modified-0.1X (data not shown) approaches produced quantifiable amounts of the target miRNA library when prepared from 60 ng mitochondrial RNA, although the total cDNA yield was approximately 7-fold higher from the modified-0.3X when compared to modified-0.1X protocol. Both modified approaches did decrease, but not prevent, adapter-dimer formation.

While 60 ng mitochondrial RNA is below the recommended library RNA input (>100 ng, NEBiolabs, Inc.), mitochondrial RNA yield from cell and tissue samples is often considerably below this threshold. Thus, it was necessary to investigate if the modified-0.3X and modified-0.1X approach could similarly produce the target miRNA library when prepared from much lower amounts of mitochondrial RNA. The modified-0.3X (Supplementary Figure 1 C, D) and modified-0.1X (data not shown) approaches both produced the target library when prepared from >1.8 ng (range 1.8-60 ng) mitochondria RNA. Total cDNA yields were, on average, 4.3-fold higher (*p*<0.05) when using the modified-0.3X protocol compared to the modified-0.1X. The presence of adapter-dimer dominated each library, regardless of the mitochondrial RNA input amount used. To combat this, all uniquely indexed libraries were combined in an equimolar pool and run across two lanes of a TBE polyacrylamide gel. The gel fragment corresponding to the miRNA region was manually excised, extracted from the gel and then sequenced. There were no differences in total RNA-seq reads (*p*=0.79) and total mapped miRNA reads (*p*=0.19) between the modified-0.3X and modified-0.1X protocols (Figure 4A). Over 200 miRNAs were detected across all but the lowest (1.8 ng, modified-0.1X) mitochondrial RNA inputs (Figure 4B). The number of miRNAs detected appeared to plateau when libraries were prepared from 15-60 ng mitochondrial RNA (Figure 4B) and suggests that RNA inputs of at least 15 ng are sufficient to detect most miRNAs within a sample. Ultimately, either modified approach is suitable for the construction of miRNA libraries. However, the significantly higher library yields make the modified-0.3X approach amenable to library pooling, gel excision, and subsequent sequencing.

**Figure 4.**
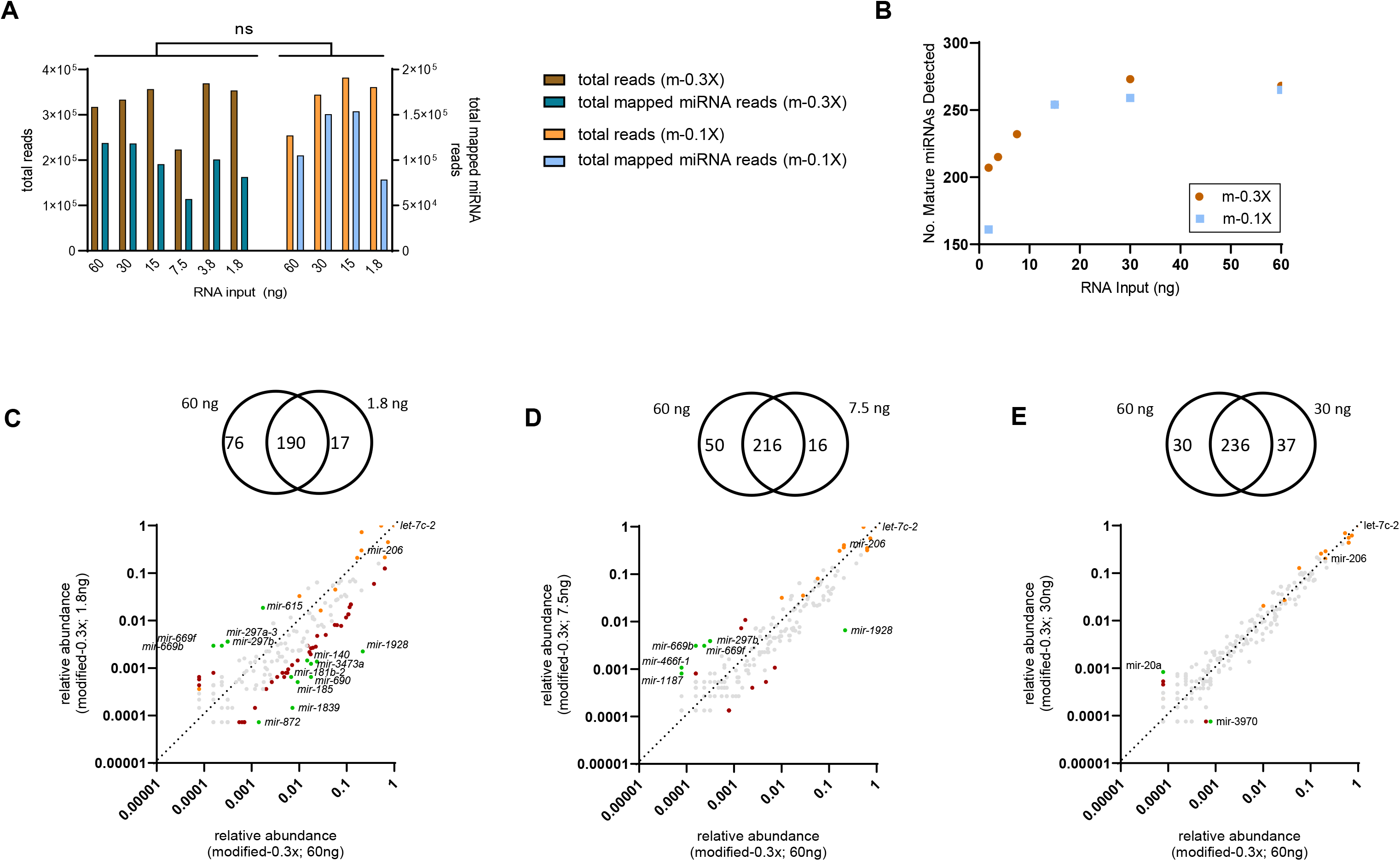
Small RNA-Seq on isolated mitochondria from L6 myocytes. **A)** Small RNA libraries were prepared from 1.8 to 60 ng mitochondria RNA with modified-0.3X (n=6) or modified-0.1X (*n*=4) protocols. The total number of reads (*p*=0.79) and total mapped miRNA reads (*p*=0.19) were not different between protocols. Student’s paired t-test, *n*=1 at each RNA input. **B)** Neither mitochondrial RNA input nor library preparation protocols limit the number of mature miRNAs that can be detected from mitochondrial RNA. The number of mature miRNA species detected across all mitochondria RNA inputs was not different (*p*=0.60, student’s paired t-test) following modified-0.3X (m-0.3X; *n*=6) and modified-0.1X (m-0.1X; *n*=4) protocols. Each point represents an independent library prepared from 1.8-60 ng mitochondrial RNA. **C-E)** The relative abundance of miRNAs within each sample is comparable between small RNA libraries prepared from high, moderate and low mitochondrial RNA inputs. Venn diagrams depicting unique and shared miRNAs detected from 60 ng mitochondria RNA when compared to **C)** 1.8 ng, **D)** 7.5 ng and **E)** 30 ng mitochondria RNA, prepared using the modified-0.3X library preparation approach. The relative distribution of miRNAs detected at each RNA input was expressed relative to miR-let-7c-2. Differences in the relative distribution of miRNAs between the highest RNA input and serial RNA dilutions are depicted by grey dots (<5-fold change when compared to 60ng), red dots (>5-fold change when compared to 60ng) and green dots (>10-fold change when compared to 60ng). Highly abundant miRNAs across all RNA inputs (including the let-7 family and mmu-miR-206) are shown in orange dots.

We next investigated the relative abundance of miRNAs sequenced from serial dilutions of mitochondrial RNA input (1.8-30 ng), when compared to the highest mitochondria RNA input (60 ng). Most miRNAs detected were common between libraries prepared from 60 ng mitochondria RNA and the lower mitochondria RNA inputs (Figure 4C-E). Further to this, the relative abundance of miRNAs was comparable between libraries prepared from the two highest RNA inputs (60 vs 30 ng; Figure 4E). A small number of miRNA species were over-(8 miRNAs) and under-represented (5 miRNAs) within libraries prepared from the highest RNA input (60 ng) when compared to lowest RNA input (1.8 ng; Figure 4C). MiRNAs detected in the highest, but not lowest, RNA input displayed low expression levels within the 60 ng sample, and together constituted a small proportion (1069 rpm) of total reads. Interestingly, the skeletal muscle-enriched miR-1, miR-133a, miR-133b and miR-16 and miR-486, predicted to have roles in the regulation of mitochondrial function (Nielsen et al., 2010, Russell et al., 2013), were detected at higher mitochondria RNA inputs (60 and 30 ng), but not lower inputs (1.8-15 ng; Figure 4). In contrast, the let-7 family as well as the skeletal muscle-enriched miR-206 were highly abundant in all samples, and together constituted 63±8% of all mapped miRNA reads. Importantly, the relative distribution of miRNAs within each sample provides adequate representation across serial dilutions of mitochondrial RNA, making it possible to accurately investigate differential miRNA expression across unique cell and tissue samples with variable mitochondrial RNA yields.

Together, these data suggest that careful optimization is required when preparing sRNA libraries from low amounts of RNA from purified mitochondria. To minimize adapter-dimer presence in sRNA libraries, gel excision and purification of the target miRNA fraction is recommended. Importantly, our data demonstrate that RNA-seq results were comparable across serial dilutions of mitochondrial RNA input, suggesting that low mitochondria RNA input does not limit the number nor the relative distribution of miRNA species that can be detected.

The transcriptomic profile of enzymatically treated isolated mitochondria from rat skeletal muscle tissue reveals a similar degree of purity as mitoplasts.

Next, we applied the purification strategies described thus far to study the transcriptome of mitochondria isolated from muscle tissue samples. From a pooled homogenate of rat skeletal muscle tissue, we compared mitochondria isolated with our IP+enzymatic treatment method against mitochondria isolated using a traditional differential centrifugation method. At the protein level, mitochondria isolated by immunoprecipitation had no detectable ribosomal marker proteins and very little abundance of peroxisome, golgi apparatus or endoplasmic reticulum marker proteins, which contrasted with mitochondria obtained via the differential centrifugation method (Supplementary Figure 2A&B). Next, to confirm the lack of co-purifying contaminants on the outer mitochondrial membrane, we compared our IP isolated+enzymatic treated mitochondria with mitoplasts (i.e. mitochondria whose OMM is removed or disrupted but with an IMM and matrix that remains intact). We generated mitoplasts and confirmed that they were devoid of outer mitochondrial membrane (OMM) proteins such as TOMM20 and inner mitochondrial membrane (IMM) and inter membrane space (IMS) proteins such as mitofilin and AIF without the loss of mitochondrial matrix proteins such as citrate synthase (Supplementary Figure 2C).

We then performed RNA-sequencing on four different mitochondrial sample types: *i*) RNase-A treated mitoplasts, *ii*) RNase-A treated IP isolated mitochondria with or *iii*) without proteinase-K enzymatic purification (to evaluate RNAs that may be bound to ribosomes on the OMM or other RNA binding proteins which protect them from RNase-A), and *iv*) RNase-A treated mitochondria isolated via differential centrifugation (“CrudeMito”). Principal component analysis of the transcriptomes of these mitochondrial sample types revealed a separation along the first principal component between crude mitochondria extracts isolated by differential centrifugation (“CrudeMito”) compared with mitochondria isolated via IP and treated with RNase-A and with or without proteinase-K treatment (MitoRNase+ProtK and MitoRNase, respectively) or mitoplasts treated with RNase-A (“MitoplastRNase”) (Figure 5A). The overlap of the latter three suggests a high overall degree of similarity of the transcriptomic profile between immunoprecipitation-based methods when compared to the crude mitochondrial fraction. For this reason, the crude mitochondrial fraction was excluded from further analysis.

**Figure 5.**
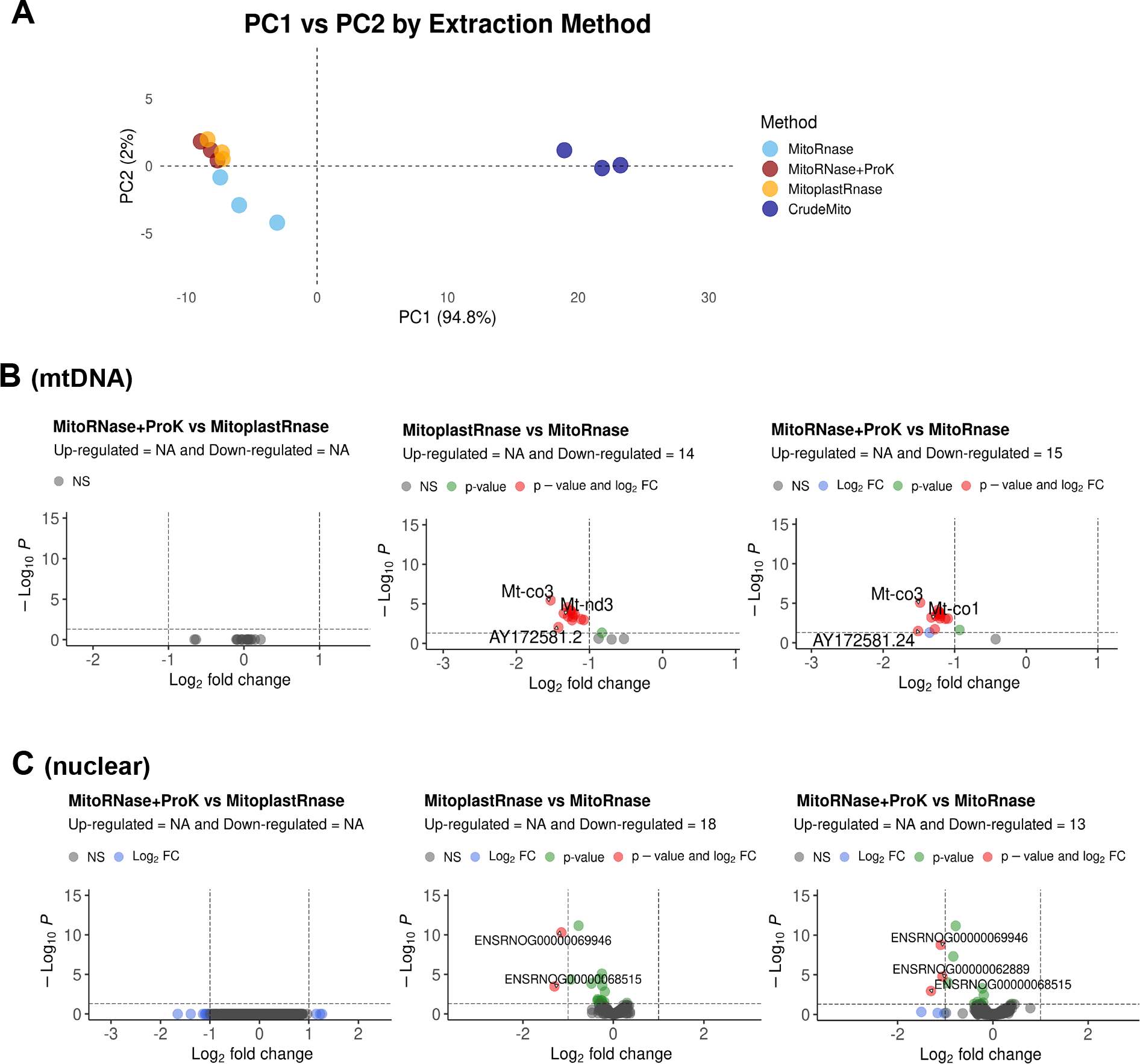
Mitoplasts and enzymatically-treated isolated mitochondria from rat skeletal muscle tissue have a similar RNA transcriptome. **A)** PCA plot of RNA-seq whole transcriptome of mitochondria isolated via IP, RNase-A treated, then treated with proteinase-K (“MitoRnase+ProtK”) or without (“MitoRnase”), mitoplast preparations treated with RNase-A (“MitoplastRnase”), and crude mitochondria extracts isolated by differential centrifugation treated with RNase-A (“CrudeMito”). Data are *N*=3 technical replicates (independent mitochondrial isolation procedures), from the tissue of *n*=1 animal. Volcano plots of differentially expressed **B)** mtDNA encoded genes, and **C)** nuclear-encoded genes between the different mitochondrial treatments and mitoplasts, statistical significance accepted at adjusted *p*-value (FDR) <0.05 and log_2_ fold-change <1.

There were no differentially expressed mtDNA-encoded genes (Figure 5B left panel) or nDNA-encoded genes between MitoplastRNase and MitoRNase+ProtK (Figure 5C left panel). In contrast, there were several DE genes (both mtDNA- and nDNA-encoded) between MitoRNase without proK and MitoplastRNase (Figure 5B & 5C middle panel), and also between MitoRNase without proK and MitoRNase+ProtK (Figure 5B & 5C right panel). The same pattern was observed for mtDNA- and nDNA-encoded rRNAs and mitochondrial tRNAs (data not shown). Collectively, these data demonstrate that mitochondria isolated via our method of immunoprecipitation followed by RNase-A and proteinase-K treatments yield a transcriptome of similar purity to that of mitoplasts.

Next, we extracted RNA from IP+purified (RNase + ProtK) mitochondria and RNA from the corresponding whole tissue from the gastrocnemius skeletal muscle of 9-week old rats (Supplementary Figure 3A & B). Whole-transcriptome RNA sequencing (Supplementary Figure 3C - E) revealed that the transcriptomes of these mitochondria had a high degree of purity as indicated firstly by a low relative proportion of nuclear-encoded mRNAs unrelated to mitochondria, such as genes associated with muscle sarcomere function i.e. myosin light (e.g. *Myl1*) and heavy (e.g. *Myh4* and *Myh7*) isoforms, α-actin (*Acta1*) and troponins (e.g. *Tnnc2*, *Tnnt3*; Figure 6A). Secondly, the mitochondrial fraction also contained a low relative abundance of nuclear-encoded OXPHOS genes (including *Cox4i1)*, as expected (Figure 6B). Thirdly, there was similar abundance of all mtDNA-encoded OXPHOS mRNA transcripts including *Mt-co3* between the whole tissue and mitochondrial fractions (Figure 6C). This serves as an important internal control as mtDNA transcripts are expected to be present in both isolated mitochondria and whole tissue in similar proportions.

**Figure 6.**
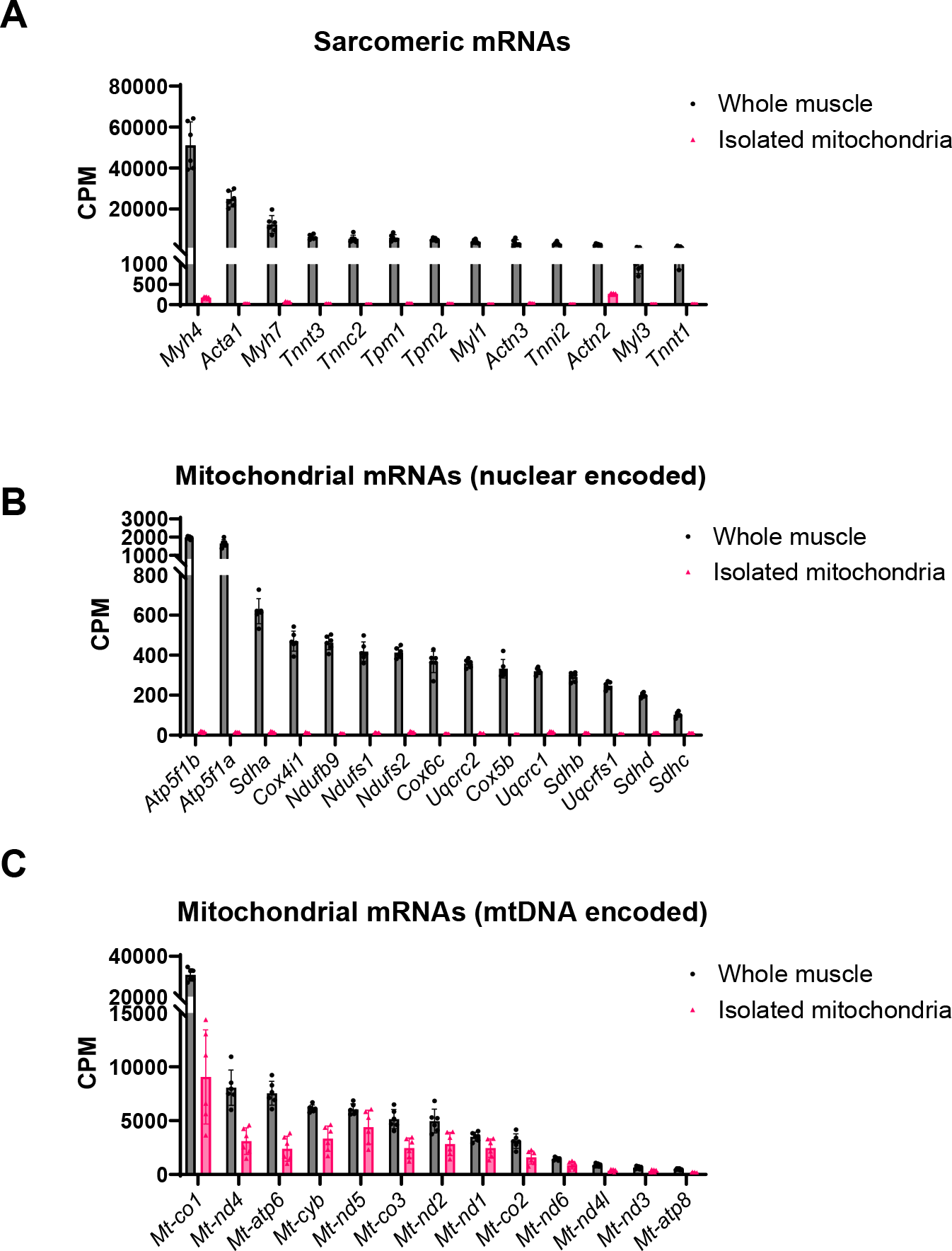
Purity of enzymatically-treated isolated mitochondria from rat skeletal muscle relative to respective whole muscle tissue. **A)** Whole-transcriptome RNA sequencing of isolated mitochondria and respective whole muscle tissue (rat gastrocnemius) shows that the isolated mitochondria were free of **D)** highly abundant muscle sarcomere related mRNAs **B)** and nuclear-encoded mRNAs involved in the mitochondrial electron transport chain, while **C)** mtDNA encoded mitochondrial electron transport chain genes were present. Each point represents the mean normalised abundance (counts per million, CPM) of an individual gene from *n*=6 animals.

Lastly, we confirmed the suitability of this approach with human muscle tissue biopsy samples. We isolated and purified mitochondria from these samples and performed analysis via RT-qPCR. This revealed the absence of *COX4I1,* used as marker of nuclear-encoded mRNA contamination, yet, several muscle-enriched miRNAs including miR-206 were detected (Supplementary Table 1), which is consistent with our results in L6 myoblasts.

## DISCUSSION

Studies of subcellular fractions using high-throughput analyses requires careful consideration of purification strategies to avoid confounding effects of contaminants. Here we report an optimised method for purification of isolated mitochondria for downstream transcriptomic analysis. Revealing the utility of this approach, we confirmed that miRNAs can be reliably detected in the mitochondria of L6 myoblasts even when libraries are prepared from RNA inputs well below the thresholds often recommended by manufacturers.

Incubation of isolated mitochondria with RNase-A digests any contaminating extra-mitochondrial RNA to ensure that only RNA species contained within the mitochondria membranes are identified in downstream analyses. This is important when investigating transcripts that do not localise in one subcellular compartment, but instead may be actively transported between compartments in response to various physiological stimuli. RNase-A protocols differ substantially between studies that have investigated mitochondrial RNA expression. In previous studies, RNase-A has been used at variable final concentrations (5-50 μg/mL) (Barrey et al., 2011, Mercer et al., 2011, Zhang et al., 2014, Bandiera et al., 2011) or has been omitted entirely (Das et al., 2012, Dasgupta et al., 2015, Lung et al., 2006, Sripada et al., 2012). Here, we demonstrate that RNase-A concentrations of at least 100 µg/mL are required to digest the majority of nuclear-encoded transcripts (such as *Cox4i1*) in rat myoblasts. Importantly, this approach is transferable between cell and tissue models and is shown to effectively degrade potentially contaminating, non-mitochondrial RNA in mitochondria isolated from human skeletal muscle tissue (Supplementary Table 1).

Investigations into the mitochondrial RNA population are often limited by technical challenges. For example, the inherently low mitochondrial RNA yields compared to total cellular RNA are well below the threshold required for commercially available small RNA library preparation protocols. This challenge may be overcome by combining mitochondria RNA from biological samples into pools that represent a single ‘intervention’ or ‘clinical’ group, and the ‘control’ group (Jagannathan et al., 2015, Wang et al., 2017). While this effectively increases the amount of mitochondrial RNA available for sRNA library preparation, this approach introduces further limitations. First, the use of pooled samples requires a much larger sample size to achieve the same statistical power (Takele Assefa et al., 2020). Second, gene expression is highly variable between individuals; taking a pooled sample approach may mask potential individual variability in miRNA expression (Takele Assefa et al., 2020, Rajkumar et al., 2015). Our findings illustrate that small RNA libraries can be prepared from individual biological samples without the need to pool multiple samples, thereby increasing the statistical power available, especially for human studies with limited sample material and high participant burden.

A modified small RNA library preparation protocol increased ligation efficiency of the 3’ and 5’ adapters to miRNAs within mitochondrial samples. However, this modified approach was not able to prevent adapter-dimer formation entirely. As the adapter-dimer product is approximately 20 bp smaller than adapter-ligated miRNAs and preferentially binds to the flow cell during RNA-Seq, gel extraction of the target miRNA region is essential to maximise read counts that map to miRNAs rather than adapter-dimer (Shore et al., 2016). When used together, the modified-0.3X protocol plus gel extraction were successful in detecting over 200 miRNAs from L6 mitochondria RNA inputs as low as 1.8 ng. Importantly, the relative distribution of miRNAs within each sample was reasonably consistent as RNA input decreased. However, some miRNA species tended to be over-represented at the lowest mitochondria RNA inputs. This may be because there were less miRNA species detected in the library prepared from 1.8 ng (207 miRNAs) when compared to 60 ng mitochondria RNA (266 miRNAs). Overall, there appears to be minimal bias towards over- or under-representing specific miRNA species at low RNA inputs.

Of note, the skeletal muscle-enriched miR-1, miR-133a and miR-133b were detected at high (30 and 60 ng) but not low (1.8-15 ng) mitochondrial RNA inputs. MiR-1 and miR-133a have previously been observed in mitochondria isolated from human skeletal and rodent cardiac muscle cells *in vitro* (Barrey et al., 2011, Jagannathan et al., 2015, Zhang et al., 2014). MiR-1 enhances the transcription of the mitochondrial-encoded *Mt-*c*o1, Mt-nd1* and *Mt-cytb*, among others (Zhang et al., 2014), while miR-133a inhibits the transcription of nuclear-encoded genes including *Ppargc1a* and *Tfam* (Nie et al., 2016), both of which are upstream regulators of mitochondrial biogenesis. The use of proliferating L6 myocytes in this study, compared with primary human myocytes (Barrey et al., 2011) and rat neonatal cardiomyocytes (Zhang et al., 2014) used previously, may partially account for the differences in miRNA species detected. Although L6 cells have reportedly higher rates of aerobic metabolism when compared to C2C12 cells and primary human myocytes, cell culture models are more dependent upon anaerobic glycolysis (Abdelmoez et al., 2020). Thus, mitochondrial abundance and function in various cell culture models may differ to those observed with *in vivo* models. This study was primarily designed to optimise the sRNA-Seq library preparation protocol rather than investigate the physiological relevance of specific miRNAs. Future *in vivo* studies are now required to profile mitochondrial miRNAs and their roles in tissues that have high bioenergetic requirements such as skeletal and cardiac muscle.

Finally, having a reliable method to discover nuclear-encoded RNAs within mitochondria in response to physiological stimuli will allow for future studies into RNA import and/or regulation at the subcellular level. For instance, non-coding RNAs such as lncRNAs can have important regulatory roles (Wadley et al., 2019), mediating cellular processes by modulating the expression of genes, stability of transcripts and/or function of proteins (Long et al., 2017), including those that directly determine mitochondrial function. However, only a relatively small fraction (∼5-10%) of all identified lncRNAs have been functionally characterized (Gao et al., 2020). Therefore, understanding the subcellular distribution may also open new avenues of investigation into how they are imported into the mitochondrial compartment, what are their putative binding target(s) and whether they modulate expression levels of mtDNA encoded genes and/or function of mitochondrial proteins. This will be key to understanding the potential effects on mitochondrial parameters such as oxidative phosphorylation, ROS production, mitochondrial biogenesis, dynamics or mitophagy.

In summary, we report a method for obtaining a highly pure fraction of mitochondria from cultured myoblasts and skeletal muscle tissue and optimised approach for high throughput transcriptomic sequencing on this subcellular fraction. Given the increasing interest in RNA-mediated regulation of cellular function in health and disease, future investigations are warranted to better understand the potential biological and physiological significance of the mitochondrial localisation of nuclear-encoded RNAs.

## MATERIALS AND METHODS

### Cell culture

L6 rat myoblast cells were obtained from a commercial vendor (ATCC Cat# CRL-1458, RRID: CVCL_0385). Cells were cultured at 37℃ with 5% CO_2_ in a humidified incubator in media consisting of Dulbecco’s Modified Eagle Medium (DMEM; Gibco #11995-065) supplemented with 10% v/v FBS (Gibco #A3161001), 100 U/ml penicillin and 100 μg/ml streptomycin (Gibco #15140-122). Cells were grown in 150 mm dishes to ∼90% confluence then washed with sterile PBS and detached (Gibco TrypLE Express #LTS12604021). Cells used for all experiments were ≤9 passages from stock and tested negative for mycoplasma contamination (Agilent MycoSensor QPCR Assay Kit, 302107).

### Myoblast mitochondrial isolation

Mitochondria were isolated from L6 myoblasts via immunoprecipitation (Miltenyi Biotek #130-096-946). Briefly, ∼5×210^6^ cells (suspended in a small volume of PBS after detachment) were transferred to 1 mL ice-cold lysis buffer on ice, then mechanically homogenised with 20 passes in a 7 mL glass dounce homogeniser (#357542, Wheaton, NJ). Wash buffer (9 mL) was added to the lysate and transferred to a falcon tube. Anti-TOMM22 antibody conjugated magnetic beads (50 µL) were added to the 10 mL lysate and incubated with gentle inversion for 1 hour at 4℃. Lysate was passed through a pre-separation filter then flowed through a column mounted on a magnetic rack. Mitochondria were washed 3 times prior to elution. Mitochondria were pelleted via centrifugation at 13,000 x g for 2 minutes at 4℃. Supernatant was then removed and the mitochondrial pellet resuspended in storage buffer to be used for subsequent enzymatic purification steps described below.

### Enzymatic purification of myoblast mitochondria

Protease treatment: To remove proteins not localised within intact mitochondria, aliquots of isolated mitochondria (1 µg total protein each) were incubated in a 50 µL reaction with or without triton-x (1% v/v) and with or without proteinase K (Qiagen #19131, final concentration 20 µg/mL) for 15 minutes at room temperature. Phenylmethylsulfonyl fluoride (20 mM) was added to inhibit proteinase-K. Samples were then stored at -20℃ until immunoblot analysis.

RNase treatment: To remove RNA not localised within intact mitochondria, in a 50 µL reaction, aliquots of isolated mitochondria (1 µg total protein each) first had 100 ng mRNA spike-in control added (StemMACS eGFP mRNA #130-101-114, Miltenyi Biotec), and were then incubated with or without triton-x (1% v/v) and with or without Rnase-A (Qiagen #19101, final concentration range 1-1000 µg/mL) and incubated on ice for 30 minutes. Trizol reagent was then added to simultaneously inactivate Rnase activity, lyse mitochondria and preserve RNA. Samples were then stored in Trizol at -20℃ until RNA isolation.

### Immunoblot analysis

Mitochondrial samples were resuspended in RIPA buffer (Millipore #20-188) containing protease inhibitors (Sigma #P8340). Protein concentration was determined by BCA assay (Pierce #23225). Samples were mixed with reducing loading buffer (4x Laemmli sample buffer with 10% 2-mercaptoethanol). Samples were loaded into a 4-15% gel (Bio-Rad #5678085) in addition to molecular weight marker (Bio-Rad #161-0373), which were then separated by electrophoresis. The stain-free gel was UV-activated for 1 minute and imaged for total protein (BioRad Chemi Doc XR+, Hercules, CA, USA) with ImageLab software (BioRad Image Lab v6). Protein was then transferred to a methanol soaked polyvinylidene difluoride (PVDF) membrane (Millipore Immobilon FL 0.45 µm #IPFL00010). The membrane was then PBS washed, then blocked for 1 hour (Li-Cor Intercept PBS blocking buffer). The membrane was then incubated with primary antibody in blocking buffer + 0.2% v/v Tween-20 overnight at 4°C. Primary antibodies used were: anti-TOMM20 (Abcam #ab186735; 1:1000), anti-Cytochrome-c (Abcam #ab133504; 1:1000), anti-Citrate Synthase (Cell Signaling Technology #14309; 1:1000), anti-α-Tubulin (Cell Signaling Technology #3873; 1:1000), anti-mitofilin (Abcam #ab110329; 1:1000), anti-AIF (Cell Signaling Technology #4642, 1:500), (anti-catalase (Abcam #ab1877, 1:500), anti-S6 RPL (Cell Signaling Technology #2217, 1:1000), anti-GM130 (BD Transduction Laboratories #610822, 1:1000), anti-SERCA1 (Cell Signaling Technology #4219, 1:1000), and anti-Lamin B1 (Cell Signaling Technology #12586; 1:1000). Membranes were then washed with PBST and incubated with secondary antibodies (Cell Signaling Technology) anti-rabbit or anti-mouse IgG Dylight® 680 nm (Cell Signaling Technology #5366S, #5470S) or 800 nm (Cell Signaling Technology #5151S or #5257S) at 1:10,000 in blocking buffer containing 0.2% Tween-20 and 0.01% SDS for 1 hour at room temperature. Fluorescent signal was then digitally acquired (Licor Odyssey® Infrared Imaging System, Lincoln, NE, USA).

### RNA isolation and quality control

Myoblast mitochondria samples stored in Trizol were thawed on ice and centrifuged at 10,000 x g for 10 min at 4℃. An aliquot of the supernatant was then used for RNA purification with on-column DNase-I treatment (Direct-Zol RNA Microprep R2060, Zymo Research). RNA concentration and purity was determined by spectrophotometry (NanoDrop 1000, Thermo Fisher Scientific) while RNA fragment size distribution was assessed by gel electrophoresis (Tapestation HS-RNA, Agilent Technologies).

### Reverse transcription and quantitative PCR (RT-qPCR)

Equal volumes of isolated RNA from the RNase-treated mitochondria samples (to account for differences in total RNA yield between samples due to RNase treatment) were reverse transcribed to first-strand cDNA in a 20 µL reaction along with no-template and no-RTase controls (Applied Biosystems High Capacity RT kit #4368814). Quantitative PCR was performed in triplicate (Agilent AriaMX G8830A) on 4 ng of cDNA in a 10 µL reaction consisting of SYBR green master mix (Applied Biosystems #4367659) with 0.3 µM forward and reverse primers (Table 1).

**Table 1.**
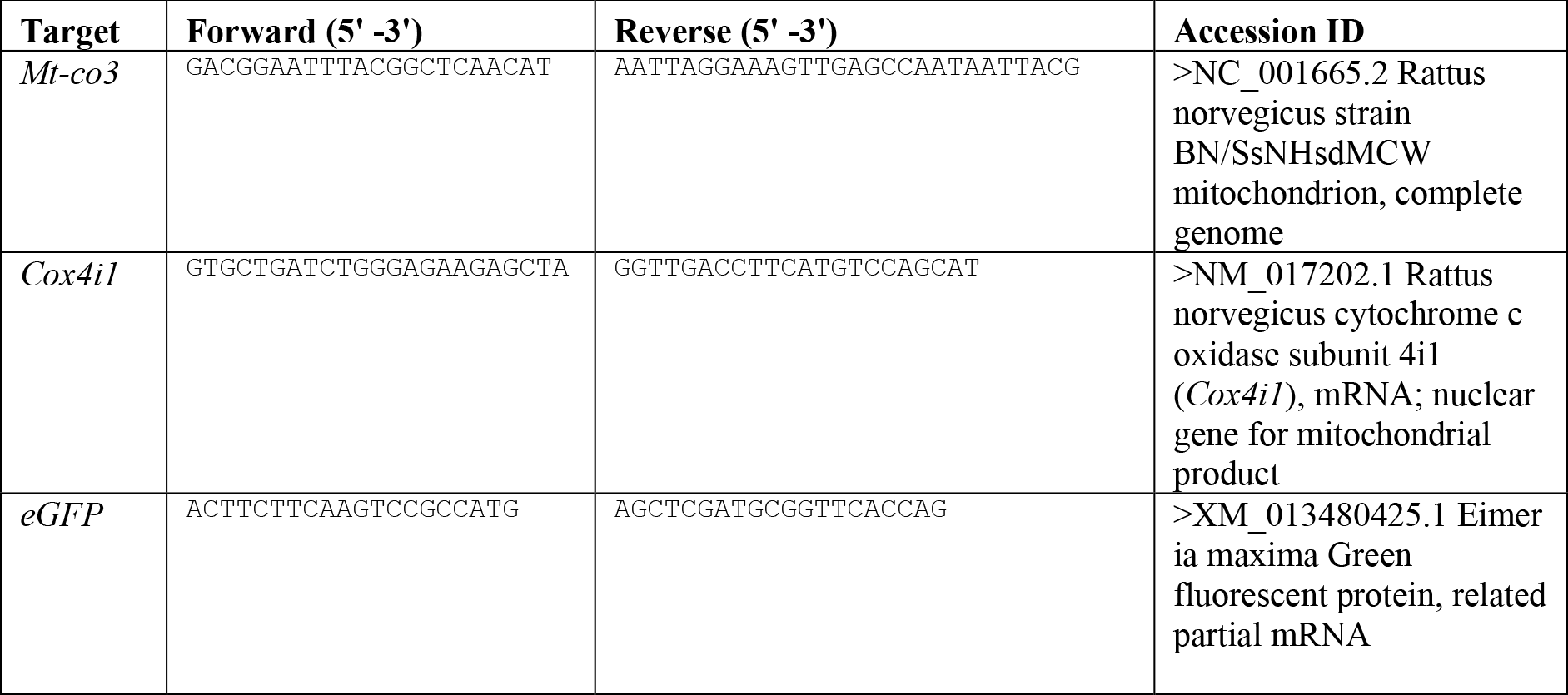
List of RT qPCR primers.

Thermal conditions used for qPCR were 10 min at 95℃ (activation) then 40 cycles of 15 s at 95°C (denature) and 60 s at 55-60℃ (anneal/extend). A dissociation curve was performed to confirm the amplification of a single product. Quantification cycle (C_q_) thresholds were calculated using software (Agilent Aria v1.5).

### Myoblast small RNA sequencing and bioinformatics

Total RNA isolated from individual (*n*=7) L6 mitochondria samples was pooled into a single tube, from which serial dilutions ranging from 0.3-20 ng/µL were prepared. cDNA libraries were then prepared using a range of mitochondrial RNA inputs (final input range 1.8-120 ng) using the NEBNext Multiplex Small RNA Library Prep Set (New England Biolabs). To investigate the efficiency of small RNA library preparation across a range of mitochondrial RNA inputs, libraries were prepared either according to the manufacturer’s recommendations for low RNA inputs (NEBNext 3’ and 5’ SR Adaptors, and RT primer for Ilumina diluted 0.5X, 15 PCR cycles) or with modifications (NEBNext 3’ and 5’ SR Adaptors, and RT primer for Ilumina diluted 0.3X or 0.1X, 20 PCR cycles). cDNA concentration was determined by fluorometric quantitation (Qubit 1xdsDNA, Thermo Fisher Scientific) while cDNA fragment size distribution was assessed by gel electrophoresis (Tapestation HS-D1000, Agilent Technologies).

An equimolar pool of all uniquely indexed small RNA libraries was prepared. Twenty-five µL of the equimolar pool was loaded across two lanes on a 6% Novex TBE polyacrylamide gel (Thermo Fischer Scientific), with 5 μL Quick-Load pBR322 DNA-MspI Molecular Marker (New England Biolabs Inc.) in a separate lane. The buffer dam was filled with 600 mL 1X TBE running buffer and the gel run at 120V for 1h. The gel was then incubated with 50 mL 1X TBE spiked with 1X SYBR Gold Nucleic Acid Gel Stain (Thermo Fischer Scientific), before being visualised on a Safe Imager 2.0 Blue-Light Transilluminator (Invitrogen) following exposure to UV-Blue (470 nm) light. Fragments corresponding to the 130-160 bp region were manually excised, before suspended and then crushed in 250 µL gel elution buffer (New England Biolabs Inc.). To increase the recovery of low concentrations of small nucleic acids, cDNA was precipitated overnight at -20°C with 0.3 M sodium acetate (pH 5.5), 4X volumes of 100% ethanol and 1 µL linear acrylamide carrier. Following overnight precipitation, the solution was centrifuged at 16,000x*g* for 60 minutes to pellet the cDNA fragment. The cDNA pellet was washed twice with 80% ethanol and resuspend in 12 µL TE buffer. Fragment size distribution of the gel-extracted cDNA pool was assessed by gel electrophoresis (Tapestation HS-D1000, Agilent Technologies).

RNA sequencing was performed by the Deakin University Genomics Core on the Illumina MiniSeq platform. A 1.3 pM loading pool was prepared with 20% Phi-X spike-in and was sequenced on a single-end 75 bp run. Sequenced reads underwent quality checks with FastQC before the adapter and reads <20nt were trimmed. Small RNAs, including miRNAs, are not well sequenced in the rat genome but are conserved between rat and mouse species. Reads were mapped to known mature mouse miRNAs accessible from miRbase v22.0. Raw read counts were normalised by the size of each individual library. Counts were visualised using GraphPad Prism (v7).

### Rodent study ethical approval and procedures

All experimental procedures were approved by the Deakin University Animal Ethics Committee (G02-2019). All animals were housed and treated in accordance with standards set by Deakin University’s Animal Welfare Committee, which complies with the ethical and governing principles outlined in the Australian code for the care and use of animals for scientific purposes. Wistar Kyoto male rats were obtained from the Animal Resource Centre, Perth, Western Australia. Rats were housed in pairs and maintained with a 12-hour light/dark cycle, constant temperature of 21 ± 2°C, and humidity levels between 40 and 70%. Rats had *ad libitum* access to standard chow diet and tap water. At 9-weeks of age, animals were humanely killed following dissection of the heart under heavy anaesthesia using 5% isoflurane gas. The gastrocnemius muscle was rapidly excised then the inner red portion was dissected. A portion of this was snap frozen in liquid nitrogen for subsequent whole muscle RNA analyses, and a separate ∼ 100 mg piece was taken for mitochondrial isolation.

### Rat muscle tissue mitochondrial isolation

Tissue was immediately placed in 1 mL ice-cold lysis buffer containing 5 µl/ml protease inhibitors (Sigma #P8340) and minced using fine scissors before mechanical homogenisation with an ice-cold Teflon-tipped glass dounce homogeniser (10 passes with rotation at 350 rpm). Intact mitochondria were then isolated using the magnetic-bead immunoprecipitation method as described above for L6 myoblasts except with the following RNase treatment conditions: RNase-A 340 µg/mL in a reaction with ∼20 µg total mitochondrial protein performed for 1 hour at 37℃, after which 5 μL proteinase-K (stock concentration 600 mAU/mL) was then added to inactivate RNase activity (Barrey et al., 2011). Each tube was briefly inverted before the mitochondria solutions were then centrifuged at 8000 x g for 10 min at 4℃. The supernatant was aspirated and the pellet was washed twice with 100 μL ice-cold storage buffer. The RNase-treated mitochondrial pellet was resuspended in 100 uL storage buffer then frozen at -80℃ until RNA extraction.

### Rat muscle tissue and isolated mitochondria RNA extraction

For RNA extraction from gastrocnemius whole muscle tissue, ∼15 mg frozen tissue was placed in a liquid nitrogen pre-chilled cyrotube with a 5 mm stainless steel bead. The tissue was then mechanically disrupted (2 cycles at 4m/s for 10 s, MP Biomedical FastPrep). Tri-reagent (600 uL) was added, then RNA extracted (Zymo DirectZol RNA Miniprep #R2050). For RNA extraction from isolated mitochondria from the respective red portion of gastrocnemius muscle, frozen samples were thawed in the presence of 5 volumes of TRI-reagent (Qiagen), pipette mixed thoroughly then centrifuged at 16,000 x g for 1 min. RNA extraction was then performed with on-column DNase I treatment (Zymo Research DirectZol RNA Microprep #2060). Extracted RNA was tested for purity on Nanodrop, concentration with Qubit HS RNA assay (Thermo Fisher) then RNA integrity number (RIN) was determined using a Tapestation (Agilent) with High Sensitivity RNA reagents. All RINs from whole tissue RNA were ≥7.0.

### Rat muscle tissue and isolated mitochondria total RNA sequencing

Total RNA from whole-tissue (50 ng) and isolated mitochondria (10 ng) were converted to cDNA libraries (Zymo-Seq RiboFree Total RNA #R300, Zymo Research). Briefly, ribosomal RNA was depleted, then remaining transcripts were fragmented and converted to cDNA using random priming. After second strand synthesis, the ends of the cDNA were enzymatically repaired and Illumina-compatible sequencing adaptors were ligated. Library size (TapeStation, Agilent) and concentration (Qubit) was assessed prior to sequencing. RNA sequencing was performed by the Deakin University Genomics Core on the Illumina NovaSeq 6000 platform. Reads underwent adapter trimming and quality check with Skewer then mapped to the rat genome (Ensembl version 99) with STAR aligner v2.7.2a. STAR generated read counts were collated in R version 4.0.3 (R Development Core Team, 2010). Normalised counts were visualised using GraphPad Prism (v7). Analysis of differential expression was performed using Voom/Limma in Degust v 4.1.5 (Powell, 2019). Transcripts with a false discovery rate (FDR) <0.05 were considered differentially expressed.

### Rat muscle tissue mitoplast preparation and RNA sequencing

All experimental procedures were approved by the Deakin University Animal Ethics Committee (G01-2023) and housed and treated in accordance with standards set by Deakin University’s Animal Welfare Committee, which complies with the ethical and governing principles outlined in the Australian code for the care and use of animals for scientific purposes. A 14 week old male Wistar Kyoto rat was obtained from the Animal Resource Centre, Perth, Western Australia and humanely killed following dissection of the heart under heavy anaesthesia using 5% isoflurane gas. The hindlimb muscles were rapidly excised and cut into 200 mg pieces. Each piece was immediately placed in 1 mL ice-cold lysis buffer containing 5ul/ml protease inhibitors (Sigma #P8340), minced using fine scissors and incubated on ice for 30 min. Samples were then mechanically homogenised with an ice-cold Teflon-tipped glass dounce homogeniser (10 passes with rotation at 350 rpm) and the homogenate was pooled. To compare different isolation methods, intact mitochondria were then isolated from separate aliquots of the same homogenate using either the magnetic-bead immunoprecipitation method described above, or a differential centrifugation method.

To obtain mitochondria by differential centrifugation, tissue homogenate was spun at 800 × g for 10 min at 4℃ and the pellet was discarded. The supernatant was then spun at 16,000 × g for 30 min at 4℃ and the pellet resuspended in 1 mL of isolation buffer containing 1mM PMSF and then spun at 10,000 × g for 10 mins at 4℃. The mitochondrial pellets were washed twice in 1 mL ice-cold storage buffer. The mitochondrial pellets were then pooled and resuspended a small volume of storage buffer.

To generate mitoplasts, intact mitochondria (40 µg) obtained from the magnetic-bead immunoprecipitation method were incubated in 0.06% digitonin on ice for 30 min, followed by 30 min incubation on ice with proteinase-K (final concentration 20 µg/mL). The samples were then incubated on ice for 10 min in PMSF (final concentration 10 mM) to inhibit proteinase-K. The mitoplast pellets were then washed once in 0.75 mL storage buffer containing 5 mM PMSF and then once in 0.75 mL storage buffer without PMSF. Each time the pellets were spun at 13,000 × g for 2 min at 4℃ and the supernatants discarded. Finally, the mitoplast pellets were resuspended in 40 µL storage buffer.

The following 4 different treatments were conducted in triplicate on 40 µg of protein from mitochondria or mitoplasts that were obtained from the hindlimb muscle of a single animal. All four treatments included 120 µg/mL RNase-A (cat# 19101, Qiagen, Hilden, Germany) incubation for 30 min on ice.

1. MitoProK: Mitochondria from magnetic-bead IP, +RNase-A (followed by 20 µg/mL proteinase-K)
2. MitoRNase: Mitochondria from magnetic-bead IP, +RNase-A (no proteinase-K)
3. MitoplastRNase: Mitoplasts, +RNase-A (no proteinase-K)
4. CrudeMito: Mitochondria from differential centrifugation, +RNase-A (no proteinase-K)

After RNase-A incubation, either proteinase-K (final concentration 20 µg/mL) or a similar volume of buffer was added to the respective treatment groups and incubated on ice for 30 min. Each tube was then centrifuged at 13,000 × g for 2 min at 4℃. The supernatant was aspirated and the pellet was washed once with 500 µL and then once with 200 µL ice-cold storage buffer. The mitochondrial pellets were resuspended in 400 µL Tri-reagent and frozen at -80℃ until RNA extraction.

For RNA extraction, frozen samples were thawed and then pipette mixed thoroughly to disrupt the pellets and then centrifuged at 13,000 x g for 10 min. RNA extraction was then performed with on-column DNase I treatment (Zymo Research DirectZol RNA Microprep #2060) and eluted in 12 µL nuclease free water. RNA yield was determined using a Tapestation (Agilent) with High Sensitivity RNA reagents. Mitochondria from magnetic-bead IP yielded ∼1.5 ng RNA per µg of protein, whereas mitochondria from differential centrifugation yielded ∼0.2 ng RNA per µg of protein.

Total RNA extracted from mitochondria from differential centrifugation (6.3 ng), mitochondria and mitoplasts isolated from magnetic-bead IP (40 ng) methods were converted to cDNA libraries (Zymo-Seq RiboFree Total RNA #R300, Zymo Research) then underwent whole transcriptome RNA sequencing as described above.

### Statistical analysis

Small RNA-seq data was analysed by paired t-test. RNase-A dose-response data were tested for normality of variance and then analysed by one- or two-way ANOVA with Tukey’s post hoc test for multiple comparisons using GraphPad Prism (v7). All data are presented as mean (SD) with significance accepted at p<0.05.

## Acknowledgments

We acknowledge Gisella Mazzarino and Amandi Gunawardena for their assistance with animal experiments and tissue collection and Erin Mayne for her assistance with the muscle tissue mitoplast preparation and RNA sequencing experiments. We acknowledge Sarah Alexander for her assistance with female participant recruitment. Severine Lamon is supported by an Australian Research Council Future Fellowship (FT10100278).

## Author contributions

Conceptualization: JLS, AJT, SN, S.Lamon, GDW

Conducted experiments: JLS, AJT, GDW, S.Loke

Conducted human studies: JLS, GDW, S.Lamon, HD

Data analysis (bioinformatics): MZ, LC, MS

Data analysis and visualisation: JLS, AJT, MS

Manuscript – original draft: JLS & AJT

Manuscript– review and editing: all authors.

Approved final version of manuscript: all authors.

## Notes

### Competing Interest Statement

The authors have declared no competing interest.

### Summary of Updates

Figures 5-7 have been merged and now appear as Figure 4. New experiments have been added as Figure 5 and supplementary Figure 2; Figure 8 has been amended and is now Figure 6; an author has been added.

