## Supplementary material for "Purification of mitochondria from skeletal muscle tissue for transcriptomic analyses reveals localisation of nuclear-encoded non-coding RNAs"

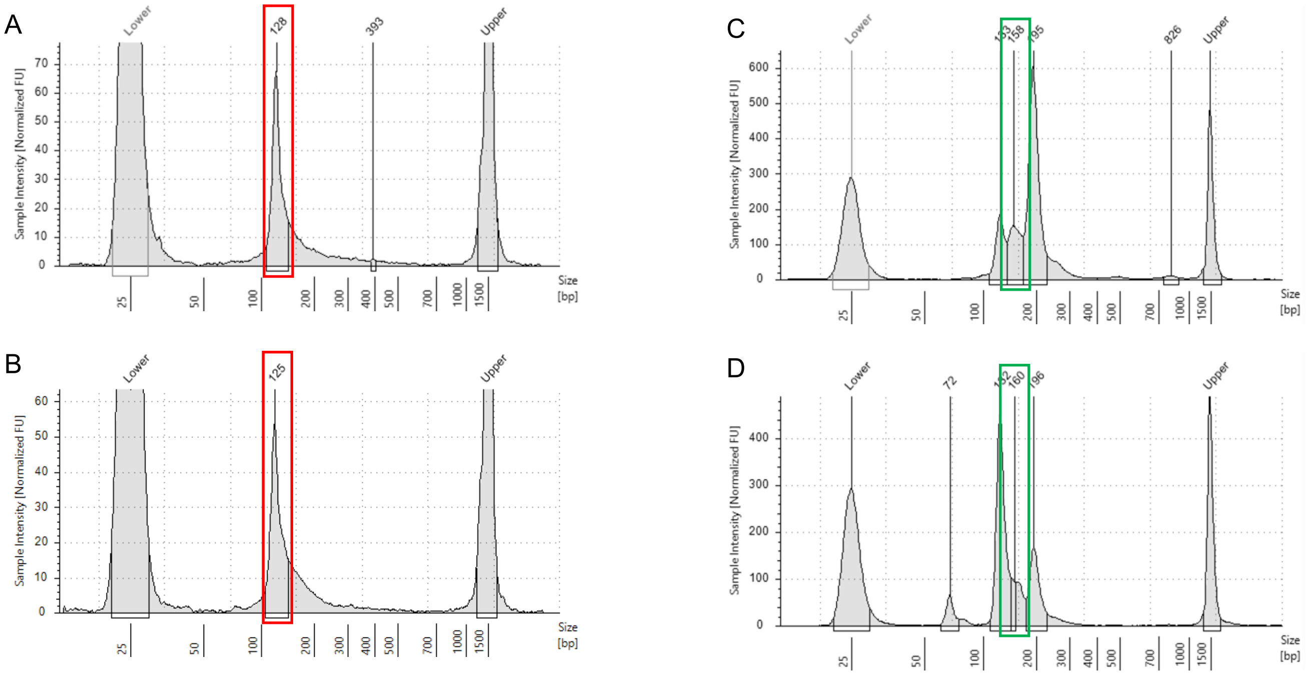


**Supplementary Figure 1. Small RNA-seq library preparation from mitochondrial RNA.** Representative electropherogram traces of libraries generated with the standard 0.5X dilution of 3’ and 5’ adapters produces large amounts of adapter-dimer (red box) and does not ligate to mitochondria miRNAs with either A) 120 ng or B) 60 ng mitochondrial RNA input. Lower molar ratios (0.3X dilution) of 3’ and 5’ adapters produces the target miRNA library (green box) with C) 60 ng and D) 1.8 ng mitochondria RNA. Peaks at 25 and 1500 bp are internal size standards.

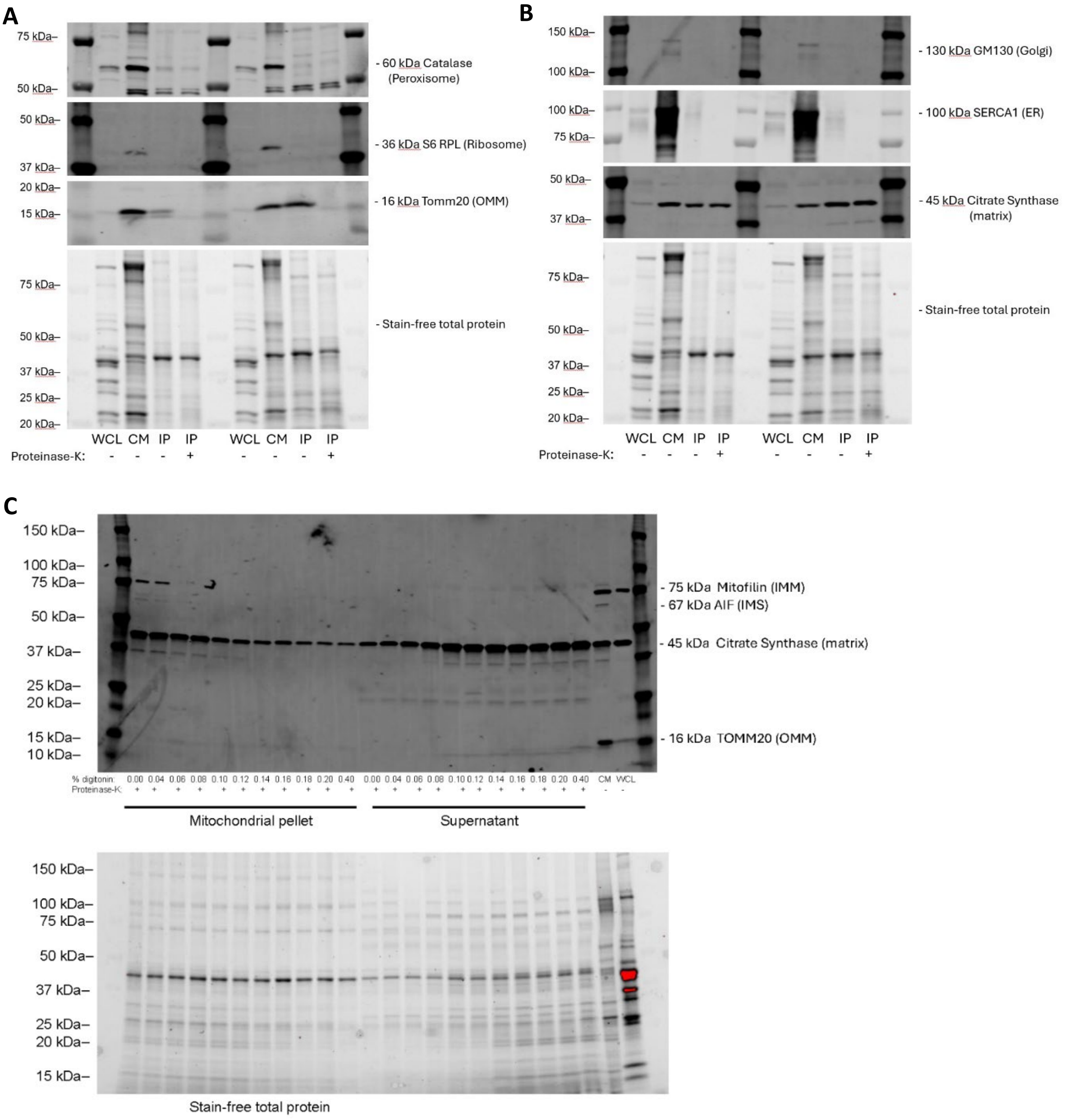


**Supplementary Figure 2. Purity of mitochondria isolated by magnetic-bead immunoprecipitation and creation of mitoplasts. A)** Immunoblots for various subcellular marker proteins of ribosomes (S6 RPL) and peroxisomes (Catalase), outer mitochondrial membrane (TOMM20) **B)** golgi apparatus (GM130) and endoplasmic reticulum (ER, SERCA1) and the mitochondrial matrix (Citrate Synthase). Samples are whole cell lysate (WCL, 6 µg total protein) along with isolated mitochondria (6 µg protein) from two independent preparations from the same rat skeletal muscle hindlimb obtained using the magnetic-bead immunoprecipitation method (IP) treated with or without 20 µg/mL proteinase-K, and differential centrifugation (crude mito, CM), as described in *Methods*. **C)** Mitoplasts created from isolated mitochondria (40 µg of protein) by incubating with increasing amounts of digitonin (0.0 – 0.40% final concentration) for 30 min at 4°C followed by 20 µg/mL proteinase-K for 30 min at 4°C. Marker proteins of subcellular compartments were then probed in the mitochondrial pellets and in the supernatants by immunoblot. Blots are representative of digitonin treatments from a single mitochondrial isolation preparation.

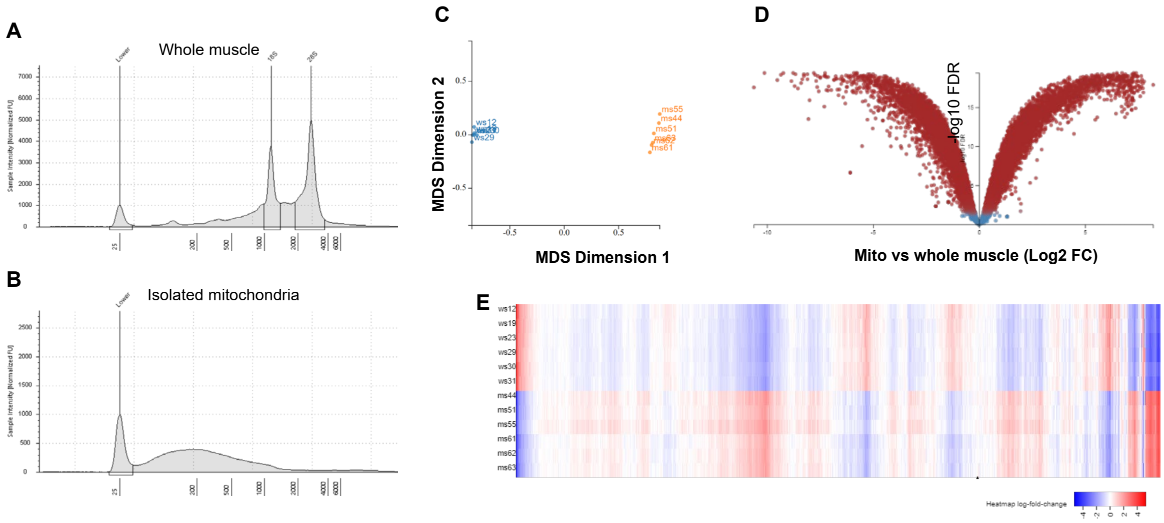


**Supplementary Figure 3. RNA from enzymatically-treated isolated mitochondria from rat skeletal muscle and respective whole muscle tissue.** Representative gel electropherogram traces of RNA extracted from **A)** whole muscle and **B)** RNaseA+ProtK treated isolated mitochondria, note the expected absence of ribosomal peaks in the mitochondrial sample. Peak at 25 nt is an internal size standard. **C)** Multidimensional scaling plot of whole muscle (blue) and mitochondrial libraries (orange). **D)** Volcano plot of genes with higher abundance in mitochondria (positive values) compared to whole muscle (red points: FDR ≤0.05). **E)** Heatmap of gene expression levels in whole muscle samples (top half) relative to the respective isolated mitochondria (bottom half) for 12,093 transcripts with ≥1 CPM from *n*=6 biological replicates.

**Supplementary Table 1: Mitochondria from human skeletal muscle contain miRNA.** Total RNA from isolated mitochondria was extracted, reverse transcribed (400 pg per reaction for mRNA analyses; 750 pg per reaction for miRNA analyses), then assessed via qPCR. RNA was free of nuclear encoded transcripts, abundant in mitochondrial genome specific transcripts, and also contained miRNAs. Quantification cycle ($C_q$) values are mean ± SD for isolated mitochondrial preparations from skeletal muscle of *n*=7 healthy female volunteers.

| **Transcript** | **Description** | **$C_q$ from mitochondrial RNA extract** |
|---|---|---|
| *MT-CO1* | Mitochondrial gene marker | 22.3 ± 2.5 |
| *MT-RNR1* | Mitochondrial 12S ribosomal RNA | 19.4 ± 2.7 |
| *MT-RNR2* | Mitochondrial 16S ribosomal RNA | 21.1 ± 2.0 |
| *COX4I1* | Nuclear gene marker | *not detected* |
| miR-1 | Muscle-enriched miRNA | 30.8 ± 0.8 |
| miR-133a | Muscle-enriched miRNA | 28.7 ± 0.8 |
| miR-133b | Muscle-enriched miRNA | 30.0 ± 0.8 |
| miR-206 | Muscle-enriched miRNA | 30.0 ± 1.7 |

### Methods for supplementary data

Human study ethical approval and procedures

All experimental procedures were approved by the Deakin University Human Research Ethics Committee (DUHREC 2014-096) and conforms to the Declaration of Helsinki. Written, informed consent was obtained from all individuals before participation. Skeletal muscle mitochondria data shown here are a subset of the female cohort (n=7; age, 23.3±3.6 y (mean±SD)) previously described by our group (Silver et al., 2020).

Skeletal muscle samples were obtained at rest from the vastus lateralis via muscle biopsy using the percutaneous muscle biopsy technique with a Bergstrom needle, modified to include suction (Bergstrom, 1962, Evans et al., 1982). Briefly, the skin was anesthetized with 1% Xylocaine, and incisions were made through the skin and muscle fascia. Approximately 60 mg freshly obtained skeletal muscle was blotted free of blood and immediately processed for the isolation of mitochondria. Each skeletal muscle sample was immediately placed in 1 mL ice-cold lysis buffer, minced and homogenised as described above for rat skeletal muscle. Intact mitochondria were then isolated using the magnetic-bead immunoprecipitation method and RNase-A treatment as described above for rat skeletal muscle mitochondria. The RNase-treated mitochondrial pellet was resuspended in 100 uL storage buffer then frozen at -80°C until RNA extraction.

Frozen mitochondrial pellets were thawed in 5X volumes of TRI-Reagent (Qiagen Inc.) and sheared through a fine pipette tip 15 times to disrupt the mitochondrial pellet. An enriched small RNA fraction was extracted from the isolated mitochondria using the miRNeasy Mini Kit with on-column DNase-I digest (Qiagen Inc.), with half-volumes of 1-bromo-3-chloropropane substituted in place of chloroform as per the manufacturer's protocol. RNA was eluted in 30 µL Nuclease-Free Water (NFW). Mitochondrial RNA concentration and fragment size was assessed by microfluidic capillary electrophoresis (2100 BioAnalyzer Small RNA Chips, Agilent Technologies).

400 pg RNA from the RNase-treated mitochondria samples was reverse transcribed to first-strand cDNA in a 20 µL reaction along with no-template and no-amplification controls (Applied Biosystems High Capacity RT kit #4368814). The RT protocol consisted of 10 min at 25˚C, 120 min at 37˚C and 5 min at 85˚C. Quantitative PCR was performed in triplicate (Agilent AriaMX G8830A). cDNA was diluted 1:5 before nuclear- (*COX4I1*) and mitochondrial-encoded (*MT-CO1, MT-RNR1* and *MT-RNR2*) transcript abundance was assessed via qPCR using Taqman hydroylsis probes Cat # 4331182 and Taqman Universal Master Mix II, no UNG (Supplementary Table 2). Thermal conditions used for qPCR were 10 min at 95°C (activation) then 40 cycles of 15 s at 95°C (denature) and 60 s at 55-60°C (anneal/extend).

**Supplementary Table 2: TaqMan hydrolysis probes used for qPCR on human mitochondria samples**

| Gene | GenBank/RefSeq | Assay ID |
|---|---|---|
| *MT-RNR1* | NC_012920.RNR1.0 | Hs02596859_g1 |
| *MT-RNR2* | NC_012920.RNR2.0 | HS02596860_s1 |
| *MT-CO1* | NC_012920.CO1.0 | Hs02596864_g1 |
| *COX4I1* | NM_001861.4 | Hs00971639_m1 |

Once the purity of the isolated human skeletal muscle mitochondria samples was confirmed, mitochondrial miRNA expression was assessed via qPCR. MiR-1, miR-133a, miR-133b and miR-206 were selected for investigation. 750 pg mitochondrial RNA was first reverse transcribed in a 20 µL reaction containing Taqman 5X primers alongside no-template controls (Applied Biosystems miRNA Reverse Transcription Kit #4366596). The RT product was diluted 1:3 in NFW before miRNA transcript abundance was assessed using Taqman miRNA hydrolysis probes (Supplementary table 3) alongside no-amplification and no-template controls using Taqman Universal Master Mix II, no UNG and Taqman 20X Assays. miRNA qPCR was performed in triplicate. Thermal conditions used for qPCR were 10 min at 95°C (activation) then 40 cycles of 15 s at 95°C (denature) and 60 s at 55-60°C (anneal/extend). Quantification cycle ($C_q$) thresholds were calculated using software (Agilent Aria v1.5) expressed after linear transformation.

**Supplementary Table 3: TaqMan miRNA hydrolysis probes used for qPCR on human mitochondria samples**

| Gene | Mature miRNA Sequence: | Assay ID |
|---|---|---|
| hsa-miR-1 | UGGAAUGUAAAGAAGUAUGUAU | 002222 |
| hsa-miR-133a | UUUGGUCCCCUUCAACCAGCUG | 002246 |
| hsa-miR-133b | UUUGGUCCCCUUCAACCAGCUA | 002247 |
| hsa-miR-206 | UGGAAUGUAAGGAAGUGUGUGG | 000510 |
